# Spatial Structure Can Decrease Symbiotic Cooperation

**DOI:** 10.1101/393868

**Authors:** Anya E. Vostinar, Charles Ofria

## Abstract

Mutualisms occur when at least two species provide a net fitness benefit to each other. These types of interactions are ubiquitous in nature, with more being discovered regularly. Mutualisms are vital to humankind: pollinators and soil microbes are critical in agriculture, bacterial microbiomes regulate our health, and domesticated animals provide us with food and companionship. Many hypotheses exist on how mutualisms evolve, however they are difficult to evaluate without bias due to the fragile and idiosyncratic systems most often investigated. Instead, we have created an artificial life simulation, Symbulation, that we use to examine mutualism evolution based on: 1) the probability of vertical transmission (symbiont being passed to offspring) and 2) the spatial structure of the environment. We found that spatial structure can lead to less mutualism at intermediate vertical transmission rates. We provide evidence that this effect is due to the ability of quasi-species to purge parasites, reducing the diversity of available symbionts. Our simulation is easily extended to test many additional hypotheses about the evolution of mutualism and serves as a general model to quantitatively compare how different environments affect the evolution of mutualism.

## 2 Introduction

Researchers have been fascinated by the existence of mutualisms – cooperation between members of different species – since Darwin [9], and therefore there are many well-documented example systems, such as pollinators with flowers, cleaner fish with other fish, and soil bacteria with plants Bshary & Grutter 7, Anstett 3, Dean et al. 10. Mutualisms between bacterial microbiomes and host species have been found to be especially common, including in many parts of the human body [4], making them relevant to human health. Further, more intricate mutualisms continue to be discovered in complex communities at an astounding rate [31, 24, 26, 23, 27]. Because these systems range across abundant sizes and species, it has been difficult to identify general principles of the evolution of mutualism. Further, most of these mutualisms are ancient, and therefore difficult to recreate or determine the factors that led to the original evolution of the behavior.

Like all cooperative systems, mutualisms are at risk of one partner cheating the other and destroying the mutually-beneficial arrangement [22]. This conflict raises the question of how mutualisms are able to arise in evolving populations and be maintained over many generations of organisms.

There are many hypotheses about what factors could influence the evolution and maintenance of mutualisms [22], several of which highlight the influence of the natural history of the partners [23], the amount of vertical and/or horizontal transmission of the symbiont [28], and the nature of the benefit conveyed by the symbiont [6]. While examining extant mutualisms can provide insight into how these factors affect their stability, it is difficult to experimentally test the influence of these factors on the *de novo* evolution and long-term maintenance of a mutualism.

The evolution of mutualisms is difficult to test in biological systems due to the long generation times of partners (even bacterial species take months to evolve [27]) and our imperfect control of the experimental system. To circumvent these challenges, we created a simulation to evolve a mutualism under varying conditions, called Symbulation. Using this system, we were able to vary specific factors while guaranteeing conditions were otherwise identical, enabling us to thoroughly test the effects of natural history, vertical transmission, benefits from the symbiont, and spatial structure on the *de novo* evolution of a mutualism.

## 3 Literature Reviewed

Mutualism – i.e. cooperation between members of different species – has been a subject of interest to field researchers since Darwin [9]. Due to this interest, there are countless examples of mutualisms in the natural world, a number of which are reviewed in depth by Herre et al. [19]. Because two organisms of different species are highly unlikely to share a cooperative gene, kin selection and the idea of selfish genes do not easily explain this type of cooperation, making it all the more intriguing, and therefore leading to an abundance of hypotheses regarding their evolution and maintenance.

Historically it was a common assumption that because mutualism was better overall for both species, parasites would be pressured to evolve into commensalists, which would then have pressure to evolve to mutualists [14]. However Ewald et al. introduced the idea that, while parasitism to mutualism is a spectrum, we must remember that selection often pressures individuals of a species to cheat a cooperative trait because in a cooperative background, cheaters will have a fitness advantage [14].

Frank et al. proposed that mutualisms are still selected due to benefits to kin, just indirectly. If the original donor is likely to have kin nearby that would benefit from reciprocation from the original recipient, it could lead to the cooperation being under more consistent positive selection [17]. Van Cleve et al. [29] modeled a system with a high benefit relative to the cost of the mutualism, as well as high within-species relatedness and high between-species partner fidelity – the idea that the maintenance of evolution depends on partners remaining with each other for extended periods of time, thus aligning their fitnesses [8]. They found that with those assumptions, feedback mechanisms between partners were more important than the genetic correlations of individuals within each of the species involved [29].

Other researchers have turned to inspiration from mate choice, drawing the parallel that a female choosing a male (or vice versa) is in many ways similar to a mutualist choosing a partner among the individuals of the other species [25]. Using game theoretic models of rational traders in human markets, assuming that partner choice is the dominant force in the system, Noe et al. found that the cost of sampling partners determined whether the host was choosy or not. If it is cheap to figure out the quality of a partner, it makes sense to try many and keep the best one. Conversely, if it expensive to try out a partner, hosts cannot switch symbionts whenever they please and so will be more tolerant of less-than-ideal partners [25]. This idea of partner-choice being a determining factor in the evolution of mutualism first arose in [8] in opposition to the concept of partner fidelity. Conversely, partner-choice is the idea discussed previously where a host has some way of sampling potential partners and picking the highest quality, thus possibly selecting for symbionts to be of higher quality [8].

Partner fidelity can also take place across generations with vertical transmission. Vertical transmission is when offspring of a host are infected by offspring of the host’s symbiont [15]. This idea of vertical transmission was first discussed in terms of parasites passed from parent to offspring, which Fine et al. modeled. However the same idea can and should be applied to potentially cooperative symbionts. In Yamamura et al., researchers presented a differential equation model of a mutualism system where vertical transmission can evolve [32]. They found with their model that when the vertical transmission rate reaches a tipping point, the system will be pushed toward perfect vertical transmission. That tipping point’s exact value can vary based on other factors, however. Bronstein et al. also found that ecological conditions must be taken into account when trying to understand the costs and benefits of mutualisms because they can cause a large shift along the parasitism-to-mutualism spectrum [5].

Much of the research on cooperation is focused on how cheaters – or lowquality partners when considering mutualisms – are prevented from invading a system [18]. However Heath et al. pointed out that, when natural systems have been closely studied, a large amount of natural variation is generally present, which many models do not take into account [18]. Therefore, when determining how mutualisms evolved and are maintained in many natural systems, models must incorporate large amounts of standing variation in partners [18]. Foster et al. found that factoring in standing variation in partner quality may actually have a stabilizing effect on mutualisms [16]. When all partners are of high quality, there is no reason to be choosy. When choosiness is no longer under strong selection, low-quality individuals can start to take advantage of non-choosy partners [16]. Therefore, if a variety of low-quality partners are introduced every generation, selection for choosiness would be maintained and therefore cooperation as well.

Yamamura et al. explored how spatial structure and dispersal could affect the evolution of a mutualism in [33]. They created a mathematical model and simulation with a mutualist and non-mutualist phenotype with a high benefitto-cost ratio and a low reproductive rate. They found that unlimited dispersal prevented the mutualist phenotype from being able to invade non-mutualists, and conversely with neighborhood reproduction only, the mutualists were able to invade non-mutualists [33]. Doebeli et al. also found that spatial structure was vital to the evolution of mutualism in their model [13]. They also required that increased investment in the mutualism led directly to increased yield and with those assumptions, mutualisms were actually relatively easy to evolve. This finding led them to suggest that, in some situations, it may not be an issue for mutualism to evolve once a symbiont is engaged with a host; it may instead be more an issue of the host’s initial defenses [13].

Akcay reviews the mechanisms discussed previously and identifies selective pathways that can lead to stability in a mutualism [1]. Beyond these mechanisms, however, Akcay comes to the conclusion that most of the existing models that he found incorporated only one of the many mechanisms listed. This finding highlights a critical flaw, as Van Cleve et al. finds that when reciprocity, genetic relatedness, and synergy (the additional fitness advantage created by the mutualism) are all included in a mathematical model, they interact in non-trivial ways and cannot be assumed to be simply additive [29]. This issue is further discussed in Hoeksema et al., which reviewed the theoretical findings of the field of mutualism evolution and noted that while many of the models work well for a specific aspect, none attempted to bring together the many factors [20]. Of course, they then point out that doing so would be difficult if not impossible in an analytical model, which is compounded by the issue that naturalists have had difficulty in determining the actual costs and benefit values in any specific system [20]. A bridge needs to be created between the theoretical and empirical researchers, and I have endeavored to begin the creation of that bridge in this work.

## 4 Methods

We created an agent-based simulation, Symbulation, that models asynchronous coevolution between host organisms and symbionts, providing a simple environment to examine the conditions under which symbionts engage in strategies that are mutualistic, at one extreme, or parasitic, at the other. Each section of this paper will have additional Methods specific to it. The following applies to all experiments.

In Symbulation, host and symbiont genomes each consist of a single number, the *resource behavior*. The resource behavior can be inclusively between −1 (antagonistic) and 1 (cooperative) and determines how the organism acts toward its partner, if it has one. At every update (time step), each host receives 25 resources, which are distributed to it and possibly its symbiont based on the host’s resource behavior value. Similarly, if the symbiont receives any resources from the host, its resource behavior value determines if it returns any resources to its host.

A resource behavior value below 0 means the organism is antagonistic toward its partner. A host with a negative resource behavior will invest that proportion of the new resources available to it into defense. A symbiont with a negative resource behavior attempts to steal resources from the host. It attacks the host based on the magnitude of its resource behavior. The amount that the symbiont’s value is more negative than the host’s value determines the proportion of resources the symbiont is able to steal from the host. For example, as shown in Fig. 1, if the host has a value of −0.3 and the symbiont a value of −0.8, the host first invests 30% of its resources into defense, meaning that those resources are no longer available for host or symbiont reproduction. 20 resources came into the host cell, 6 have now been used up and 14 remain for reproduction. The symbiont then attacks with −0.8 and because that value is negative and of greater magnitude than the host’s defense value, the symbiont is able to overcome the host defense. The symbiont is therefore able to steal 50% of the *remaining* resources. Therefore the symbiont is able to steal 7 resources and apply them to its own reproductive progress. The host then has 7 remaining resources that it is able to use for its own reproduction.

**Figure 1:**
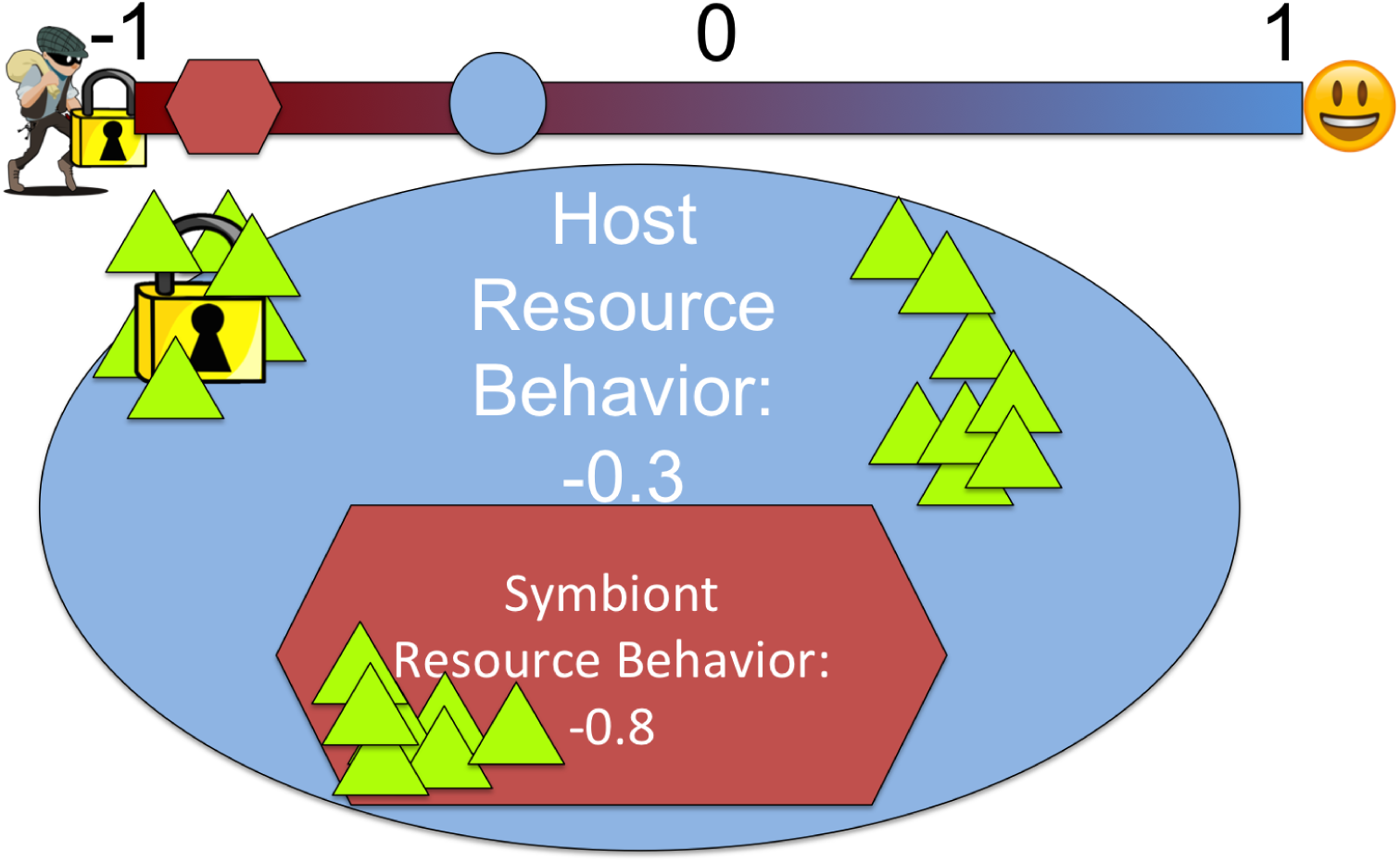
Example host and symbiont with antagonistic behavior. The host spends 30% of resources on defense (lock), the symbiont steals 50% of what remains, and the host retains what the symbiont does not steal.

Conversely, a resource behavior value above 0 means the organism is acting cooperatively toward its partner. The host donates that proportion of resources to its symbiont (if it has one), and the symbiont, in turn, donates a proportion back to the host – determined by the symbiont’s resource behavior – with a synergy factor of multiplying by five applied. This synergy factor is an abstraction of the benefits gained in a mutualistic system by dividing labor between host and symbiont. When this artificial synergy factor is replaced with two resources that organisms must specialize to use, the results are the same [30]. All combinations of positive and negative phenotypes are presented in Table 1.

**Table 1:**
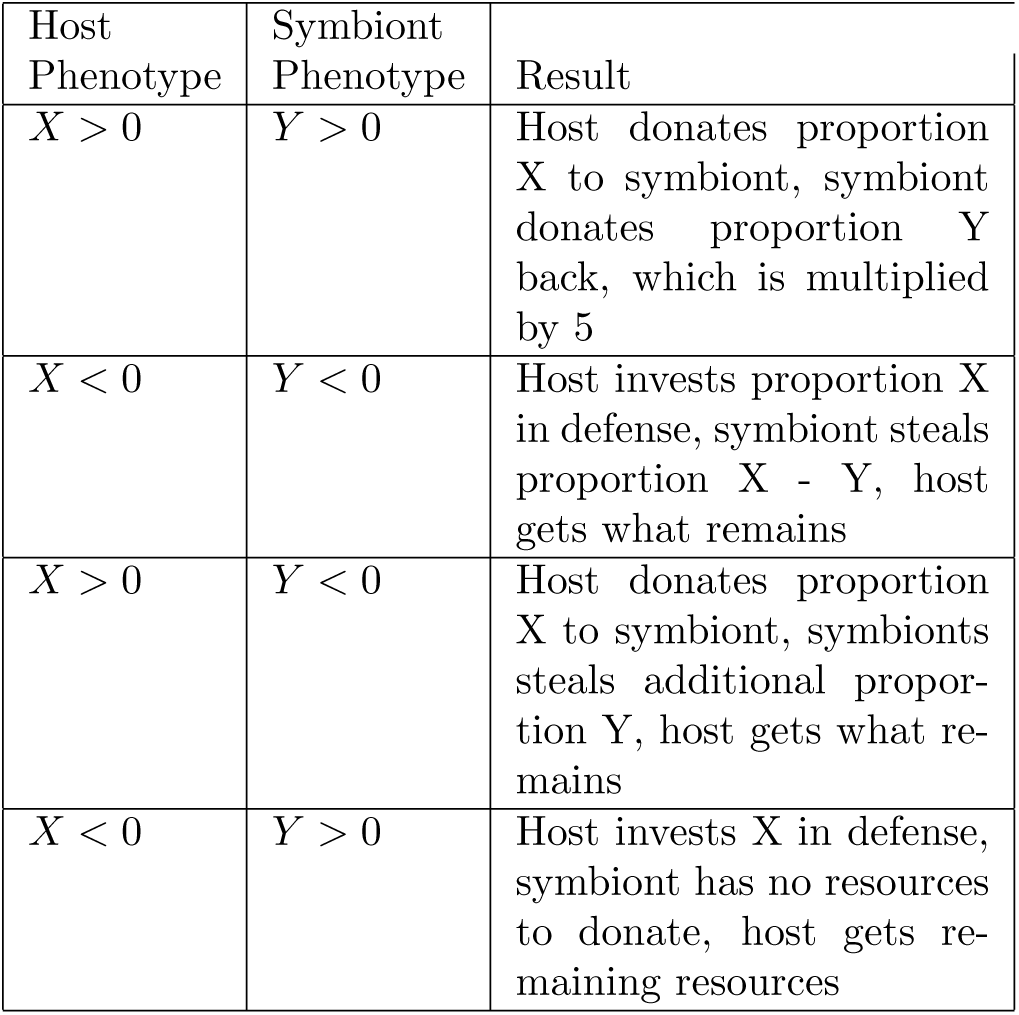
Interaction results for host and symbiont phenotypes

When a host reaches 100 stored resources, it reproduces and the offspring is placed in a random location in the world. Reproduction is imperfect, of course, and the parent and offspring both have the chance of mutation changing their resource behavior value. The resource behavior is altered by a random number pulled from a Gaussian distribution with a mean of zero and a standard deviation of 0.002. When a host offspring is placed, any pre-existing organism in that space is killed. If the host has a symbiont when it reproduces, that symbiont has a chance to be replicated and copied into the host offspring. Whether the symbiont is replicated and vertically transmitted is determined by the user-set *vertical transmission*. If the symbiont is not vertically transmitted, the host offspring is placed uninfected and keeps all of its resources every update. After reproduction, the offspring and parent both have 0 resources.

In addition to vertical transmission, a symbiont may also horizontally transmit if it accrues 50 stored resources. The symbiont offspring produced must infect a host to survive. A host in the population is randomly selected and if the host targeted is already infected, the offspring fails and dies. Hosts are limited to having one symbiont in them and once that symbiont has infected them, the symbiont may not be removed.

Determining whether a mutualism has actually emerged in any system is difficult, both because it is difficult to measure the required values and because the literature is not agreed on what the measurements or values should actually be [22]. Because Symbulation allows complete control of data, we are able to avoid any indirect proxies and instead measure exactly the behavior each partner exhibits to the other, that is, the resource behavior value. However, there is still the question of how positive a value needs to be for a mutualism and which partner needs to be positive. We are using the definition of a mutualism that the host is better off with its partner than it would be without (and conversely worse with a parasite than without). Therefore, a symbiont that has a positive resource value less than 1 (100% of resources back to host) could still be a mutualist if what it is contributing is more than the host would have without that partner, even if there are also better partners in the world. We consider a mutualism to be any relationship that appears stable over evolutionary time and both partners have positive resource behavior value. While this is not a direct test of whether the host is better off with the symbiont, selective pressure will select for hosts with a positive resource behavior value only if donating to the symbiont is of value to the host. Similarly, because defense is costly to the host, a host will only have a negative value if the symbiont is causing it more harm than not having a partner; parasitism will therefore be a stable relationship where both partners have negative resource behavior values.

Symbulation currently only supports endosymbionts, i.e. symbionts that live inside of the host and are therefore isolated from the host’s environment. All experiments had 20 replicates, ran for 100,000 updates, and had a population limit of 10,000. The default environment is a 2D well-mixed torus. All experiments start with a full population of hosts and symbionts with randomly generated interaction rate values. All code and analysis is available at https://github.com/anyaevostinar/mutualism_model/tree/master/Symbulation2.5

## 5 One Infinite Resource

### 5.1 Methods

Each section of this paper will have a methods subsection describing what additional features we made to examine the questions posed in that section. One infinite resource is the default environment and therefore has nothing unique to it. All future types of resource will have these settings unless otherwise specified in the respective methods subsection.

### 5.2 How does vertical transmission rate affect the maintenance of mutualism?

Vertical transmission is imperfect in natural systems, and the probability of vertical transmission succeeding is likely to affect the investment a symbiont places into a mutualism [22]. The more a symbiont can rely on spreading its offspring when its host reproduces, the more beneficial it would be to increase its host’s fitness. However, if it is unlikely a symbiont will be able to spread when its host reproduces, it would make more sense for the symbiont to spread via horizontal transmission [28]. Therefore, we first focused on determining how the probability of vertical transmission would influence the evolution of a mutualism. We tested values of 0, 10, 20, 30, 40, 50, 60, 70, 80, 90, and 100% vertical transmission. These set probabilities are in addition to the other requirements for vertical and horizontal transmission discussed previously. An artificial synergy factor of five was applied.

As shown in Figure 2, we found that a vertical transmission rate of 0% led to a clear parasitism where the symbiont is on average attacking the host as much as it can and the host is investing half of its resource into defense. This confirms our hypothesis that when there is no chance of the symbiont being vertically transmitted, it will not cooperate with the host and instead steal as many resources as possible for horizontal transmission.

**Figure 2:**
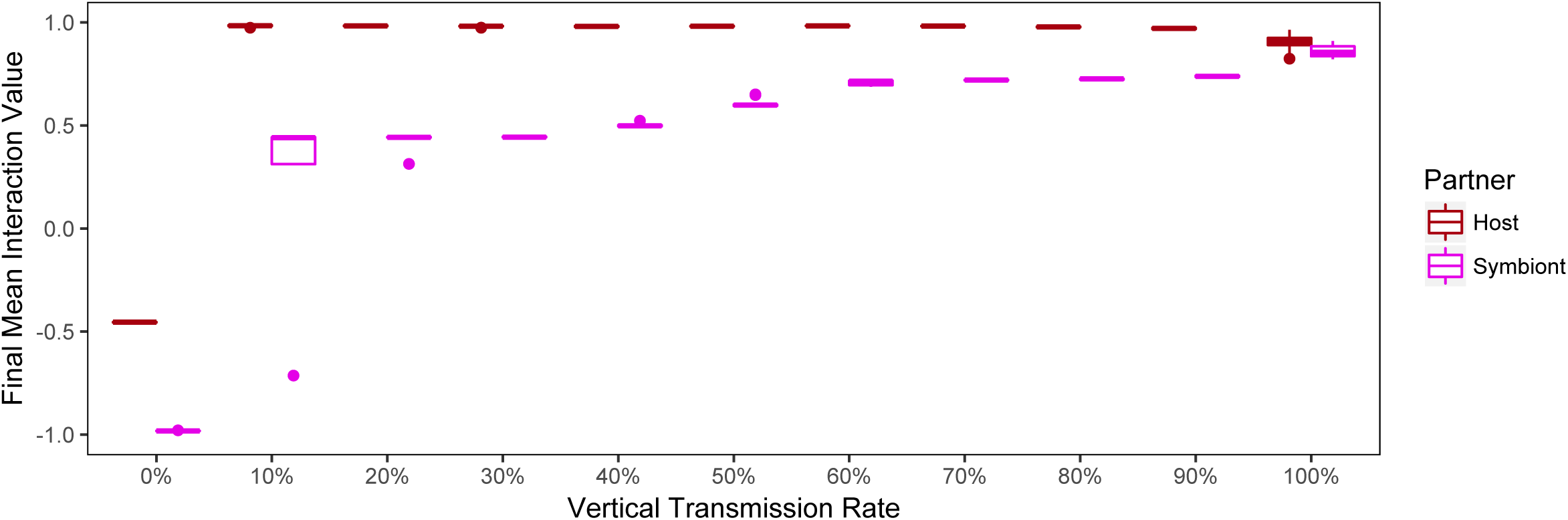
Evolution of mutualisms across vertical transmission rates. Final resource behavior values are shown after evolution across vertical transmission rates from 0% to 100%. Vertical transmission rate clearly strongly influences the final resource behavior value of each partner. At 0% vertical transmission rate, the symbiont becomes fully parasitic and the host defensive. At 100% vertical transmission rate, the symbiont and host are fully invested in a mutualism, to the point that the symbiont has lost the ability to reproduce horizontally.

Conversely, at 100% vertical transmission rate, an extreme mutualism evolves where the host donates all of its resources to the symbiont and the symbiont donates all of its resources back to the host. This result means that the symbiont has lost the ability to horizontally transfer because it can no longer amass any resources to do so. Similarly, the host is completely dependent on its symbiont such that it would be unable to reproduce without it (assuming the resources it tried to donate were therefore wasted). This extreme result confirms the hypothesis that a guaranteed vertical transmission will align the symbiont’s selective pressure with the host’s and possibly lead to a major transition over evolutionary time.

At intermediate vertical transmission rates of 10-90%, the final relationship supports the idea of imperfect mutualisms [22]. In these cases, the symbiont is not a perfect partner, but it could contribute more resources back to the host instead of keeping some for itself. However, the host still has an resource behavior value near to 1. This result is an example of when the host is still selected to invest in the mutualism with an imperfect partner because having an imperfect partner is better than no partner at all. Because of the synergy effect (x5), the host only needs to get 10% of its contributed resources back to break even. Any higher resource behavior value above 10% in a symbiont is beneficial to the host.

The transition from most likely to evolve to a mutualism and most likely to evolve to a parasitism occurs between 10 and 0% vertical transmission rate. To better understand the dynamic of this transition, we ran additional experiments at every whole percentage vertical transmission rate between 0 and 20%. In Fig. 3 it is clear that the transition between effects of vertical transmission rates are sharp between 7% and 11% with intermediate final population states at 8, 9, and 10% vertical transmission rates. This result suggests that there are few vertical transmission rates that can lead to an intermediate population state without other confounding factors, but it is possible.

**Figure 3:**
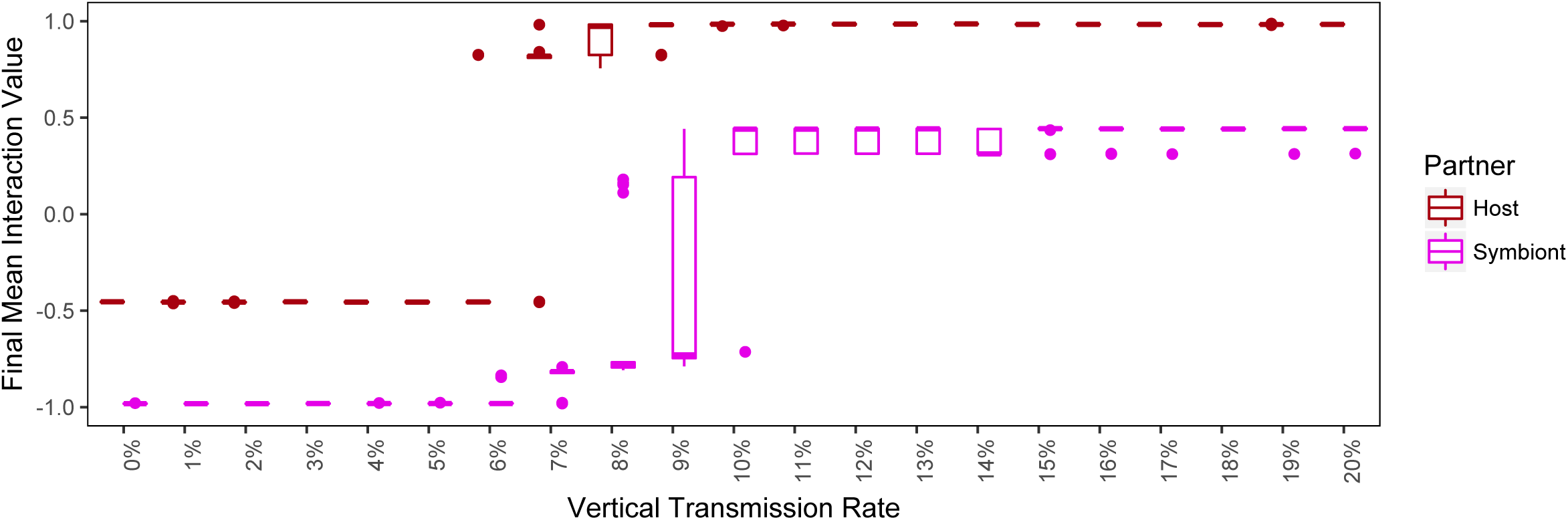
Evolution of behavior between 0% and 20% vertical transmission rates. Vertical transmission rates of 8, 9 and 10% show an intermediate final state. However, the other treatments make it clear that generally there is a sharp tipping point between a population ending in mutualism or parasitism.

The intermediate state of populations at 8 and 9% vertical transmission rate is particularly strange because the average is −0.598 and −0.286, respectively, suggesting that symbionts in these populations are at a behavior of below 0 yet the hosts are still investing in these useless symbionts. However, when the phenotypes of the symbionts are not reduced to a single average, as in Fig. 4, it is clear that these are not populations of purely parasitic symbionts and mutualistic hosts. Instead, there appears to be a stable coexistence of parasitic and mutualistic symbionts.

**Figure 4:**
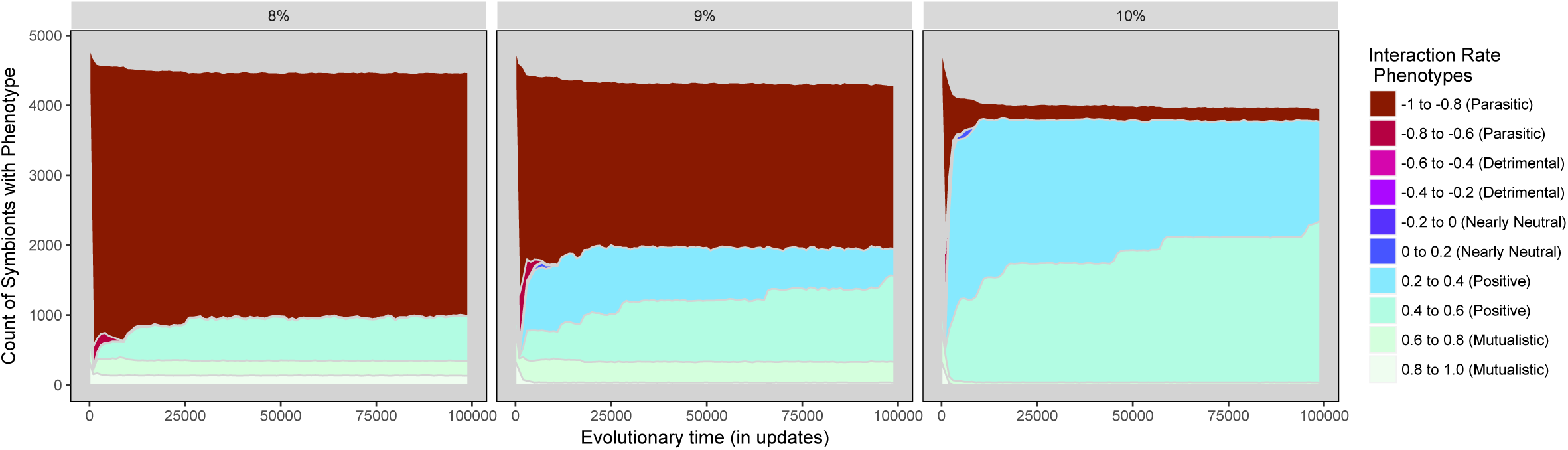
Phenotypes of symbiont at vertical transmission rates 8, 9, and 10%. At each intermediate vertical transmission rate a stable coexistence between parasitic and mutualistic symbionts persists through evolutionary time.

However, when the phenotypes of individual populations are examined, as in Fig. 5, the three possible scenarios become clear. In 17/20 replicates, a coexistence between extreme parasites and moderate to extreme mutualists was able to persist to 100,000 updates. Notably in the 3/20 replicates that did not have a stable coexistence, a mutualistic phenotype was dominant.

**Figure 5:**
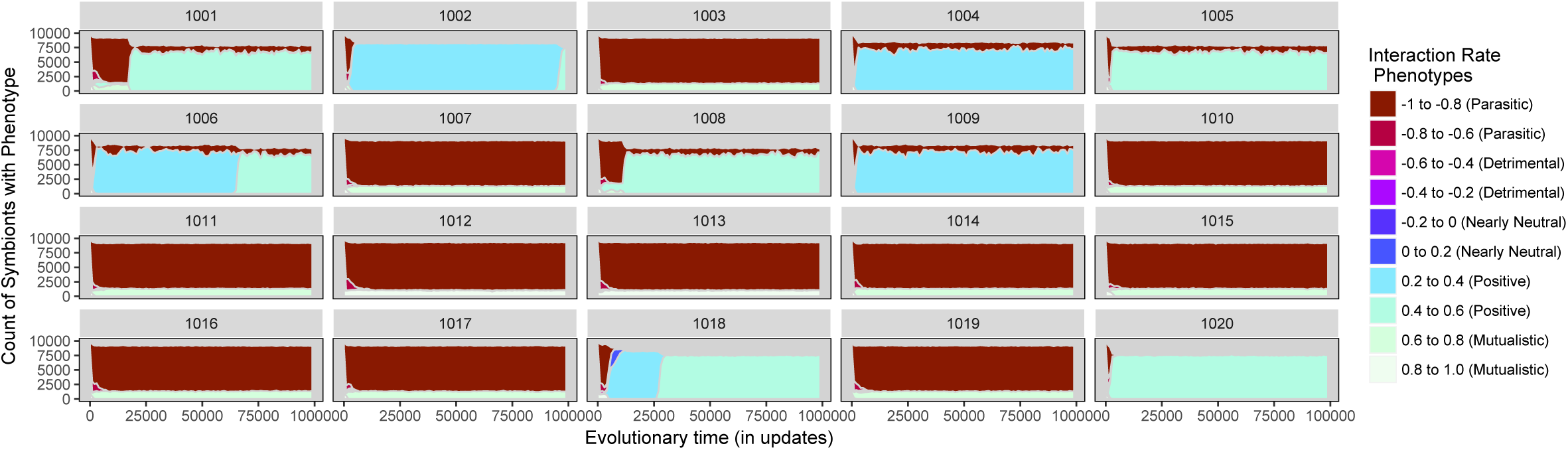
Phenotypes of each replicate symbiont population at vertical transmission rate 9%. Twenty replicate populations evolved at 9% vertical transmission rate are shown. Two stable types of co-existence exist with either imperfect mutualists or extreme parasites as dominant.

Further, there are two distinct types of coexisting populations: 1) majority moderate mutualists and minority parasites (6/20 replicates, ex: 1001 in 5) and 2) majority parasites and minority extreme mutualists (11/20 replicates, ex: 1003 in 5).

Together these results show that even in a simple agent-based system, complex dynamics can emerge and there are multiple stable states that populations can evolve to. We have confirmed that increased vertical transmission rate does more strongly select for mutualism, however at intermediate vertical transmission rates, stable coexistence of parasites and mutualists is possible.

### 5.3 How does vertical transmission affect the evolution of mutualism?

The previous section focused on whether mutualistic traits could invade an environment when starting as an established subset of the population. However, a population with high standing variation in a mutualistic trait will not necessarily have the same dynamics as a population starting with a neutral phenotype (neither mutualistic nor antagonistic) and therefore the *de novo* evolution of a mutualistic trait against a neutral background must be explored separately.

We created starting populations where all organisms started with a phenotype of 0, meaning they neither donated to their partners nor acted antagonistically. As seen in Fig. 6, at all vertical transmission rates except for 100%, starting with a neutral background led to the extinction of the symbionts before they were able to evolve to parasitism or mutualism. At 100% vertical transmission rate, a small proportion of more mutualistic hosts starts to emerge at the end of the experiment, as seen in Fig. 7, however these final populations are clearly less mutualistic than populations that start with standing diversity as discussed in the previous section.

**Figure 6:**
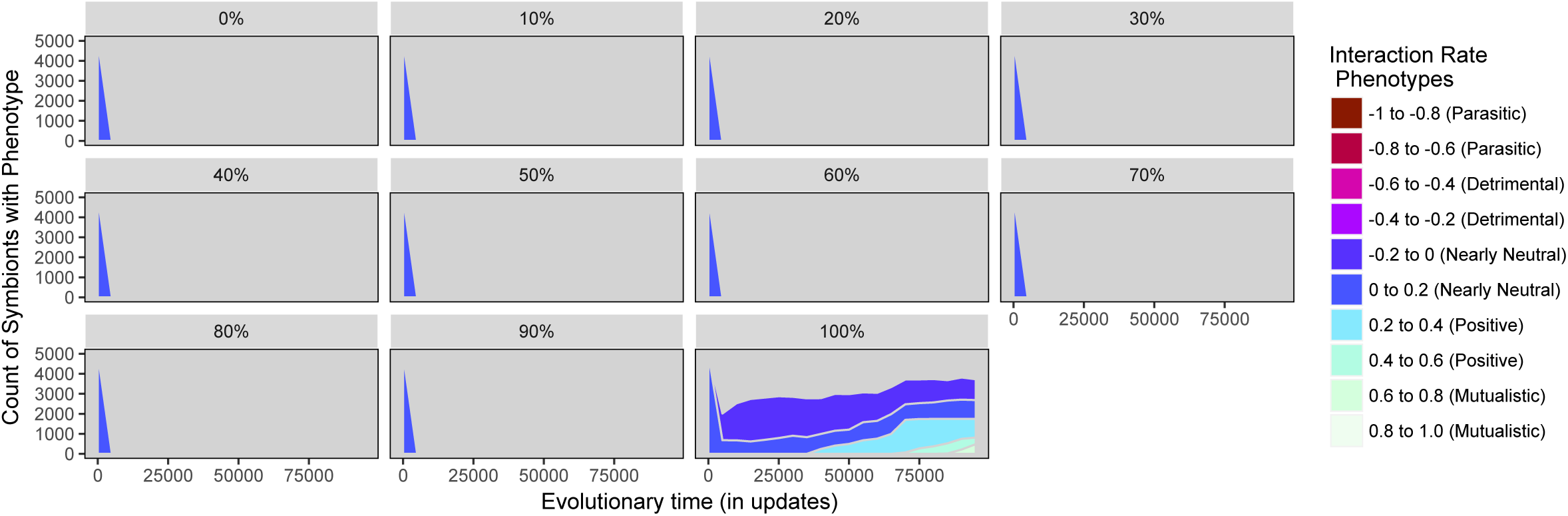
Phenotypes of Symbionts across vertical transmission rates when hosts start at neutral phenotype. Symbionts go extinct except when the vertical transmission rate is 100%.

**Figure 7:**
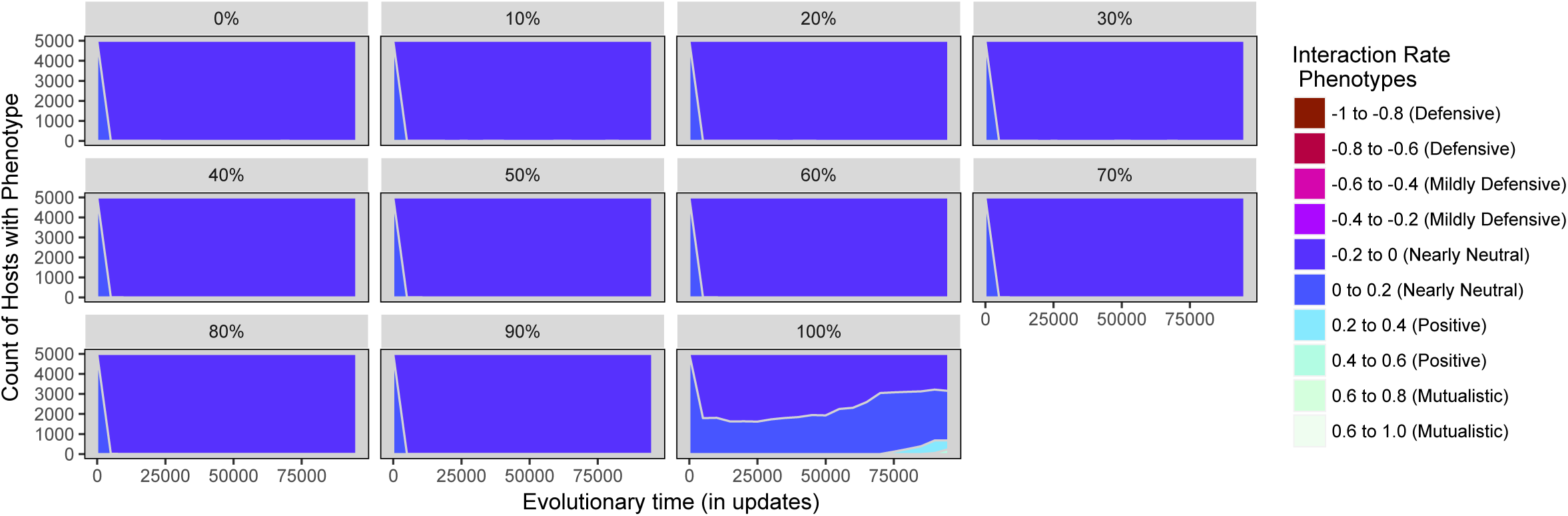
Phenotypes of hosts across vertical transmission rates when ancestral hosts all start with the neutral phenotype. Because symbionts go extinct (see 6), host resource behavior value is not under selective pressure and does not evolve except at 100% vertical transmission rate.

These results suggest that the conditions that allow mutualisms to become established and stable in a population do not necessarily also enable a mutualism to emerge *de novo* in a population of neutral phenotypes. This finding implies that other mechanisms would need to be in place to allow a mutualism to grow to a sufficient proportion of the population or allow a population to already have a diverse set of organisms when selection on the mutualist trait becomes active through exaptation.

### 5.4 How does the ancestral symbiont behavior affect the evolution and maintenance of mutualism?

The initial behaviors of symbionts is likely to influence the likelihood that mutualisms successfully evolve or can be maintained in the system. Symbionts frequently evolve to infect a new host species [22], thus beginning their coevolution with that host as potentially already parasitic or mutualistic. If a symbiont at the beginning of an evolutionary experiment has more parasitic behaviors, we hypothesized that a stable mutualistic relationship would be less likely to arise in the longer term than if the symbiont has initially random or mutualistic behaviors. Therefore, we designed three treatments to test how the symbionts’ behavior at the start of evolution changes the ending relationship between symbiont and host. Our control treatment and the default is for a new symbiont’s genome to be randomly assigned values at the start of evolution. We then created a parasitic population of symbionts to start the parasitic behavior treatment by forcing the symbionts’ resource behavior values to be negative. Conversely, we created ancestor symbionts with more mutualistic behavior by changing their resource behavior values to be positive, thus exploring the maintenance of mutualisms. The values could still be between −1 and 0 or 0 and 1, respectively.

As seen in Fig. 8 the general starting phenotype of the symbiont significantly affects whether a mutualism will evolve or be maintained at vertical transmission rates between 20% and 90%. This result indicates that the evolutionary history of the symbiont when it moves to a new host can completely alter the relationship with that host.

**Figure 8:**
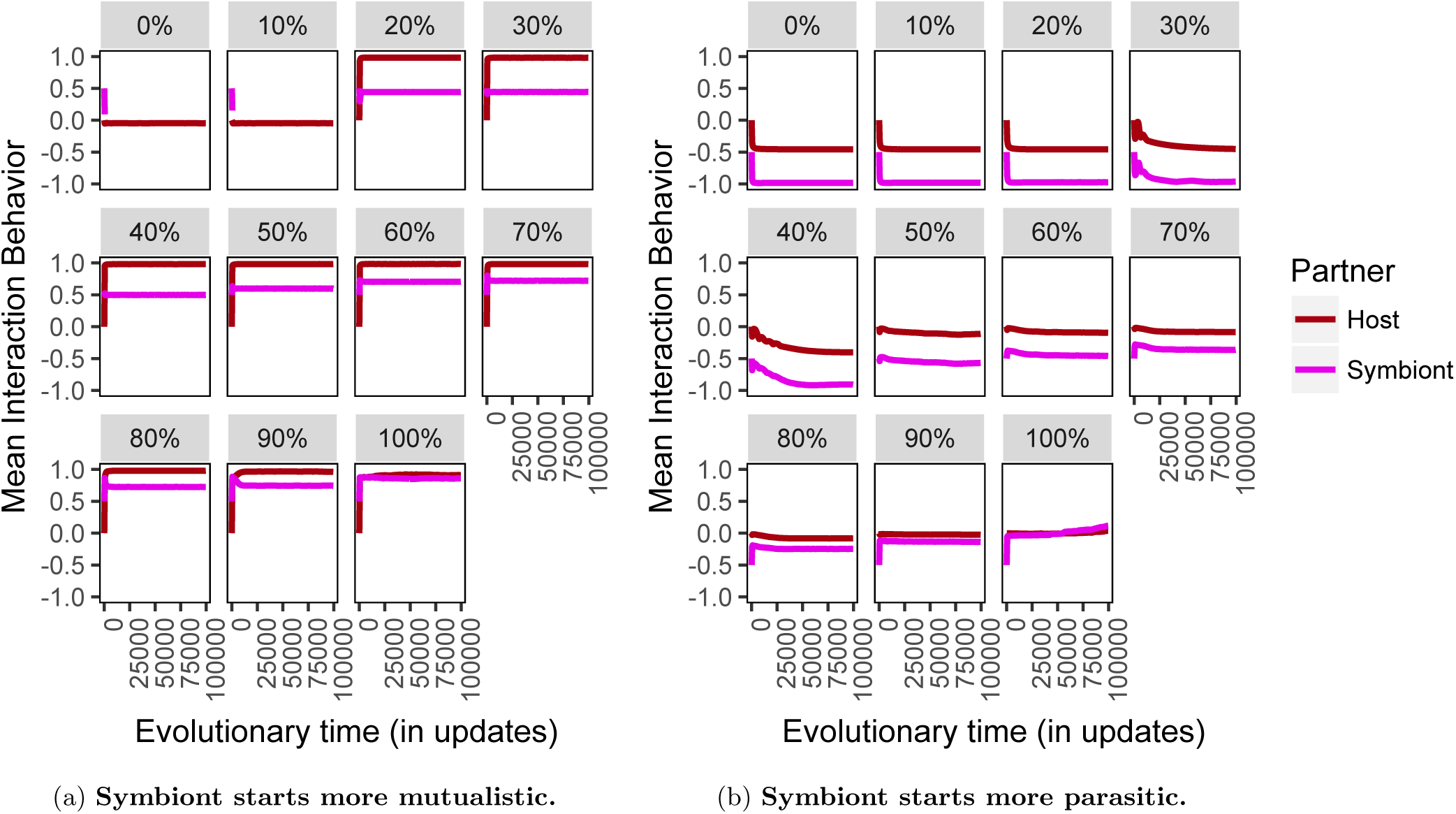
Evolution of mutualisms across vertical transmission rates when symbiont starts (a) more mutualistic than random or (b) more parasitic than random. (a) When a symbiont starts as more mutualistic in behavior, mutualisms are able to evolve at vertical transmission rates 20% and higher, where tended to fail when the symbiont started with a random behavior. (b) When a symbiont starts as more parasitic in behavior, it is more difficult for a mutualism to evolve than when the symbionts start with random behaviors. However at vertical transmission rate 100% the mutualism is able to recover from the parasitic starting behavior.

Once again the population averages in the intermediate vertical transmission rates of the parasitic starting treatments are near 0%, indicating that there is a coexistence or multiple stable phenotypes. In Fig. 9 a coexistence between two parasitic symbionts occurs at the vertical transmission rates where a mutualism would evolve if the symbiont had started with random phenotypes. This result indicates that this type of coexistence between one extreme phenotype and one less extreme phenotype is not dependent on a mutualistic host.

**Figure 9:**
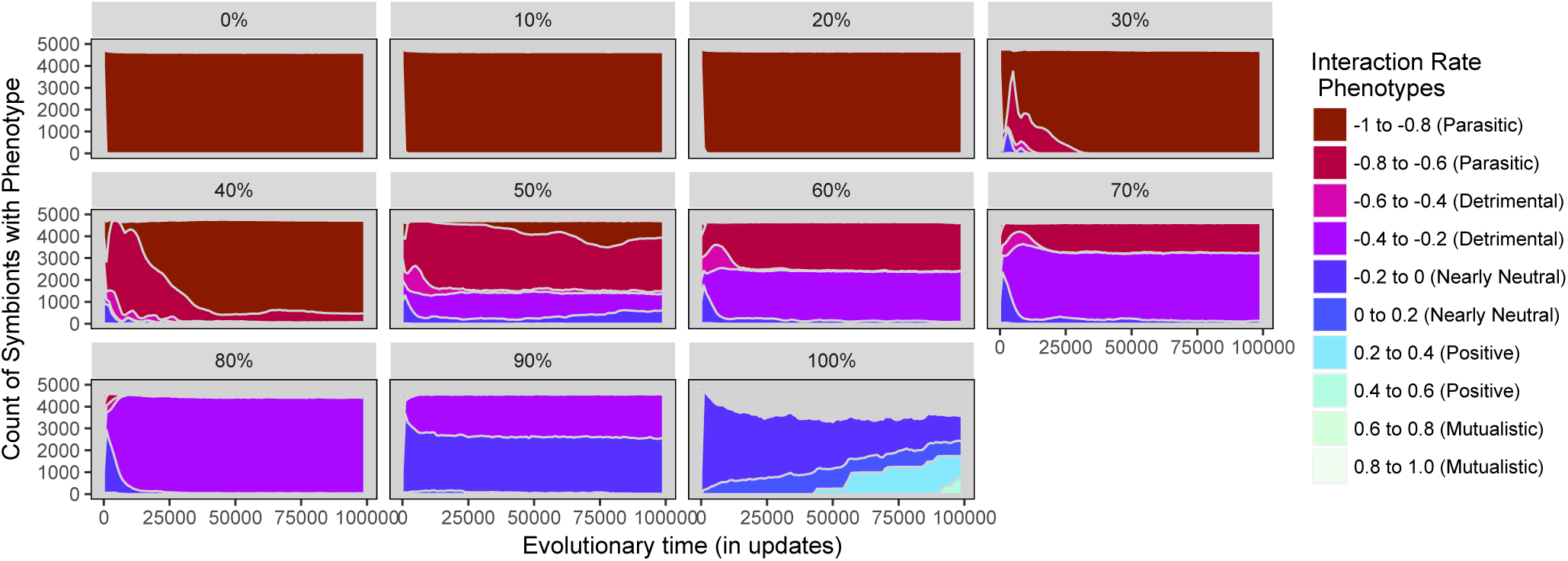
Phenotypes of symbionts starting more parasitic than random across vertical transmission rates. When starting at more parasitic phenotypes, stable coexistence of parasitic and mutualistic phenotypes can evolve only at 90% or 100% vertical transmission rate as opposed to all rates about 10% when starting random.

While the previous section shows that a mutualism cannot evolve *de novo* in this system from a neutral starting population, these results indicate that a new mutualism could evolve from a host range change of a symbiont. If the symbiont starts interacting with a new host and is more mutualistic than random, a mutualism can evolve under a broader range of vertical transmission rates.

## 6 Multiple Infinite Resources

Many natural mutualisms are successful because the endosymbiont performs a task, such as defense, that the host is incapable of doing itself (as in the classic example of the ants and acacia trees [21]). This division of labor leads to a synergy effect where the symbiont is able to increase the energy yield for the host when the host donates resources to it. In the previous experiments this effect was artificially set. Here we introduce resource types and allow the synergy effect to emerge due to true division of labor between host and symbiont. We then compare this to a control where the host is able to choose all resource types and therefore has no need of the symbiont.

### 6.1 Methods

If resource types are enabled, at least two types of resources are available in the environment in unlimited quantities, but only a set amount of each resource can be brought into the host in a given time step. If the symbiont chooses different resources than the host, its host is able to process that additional type of resource efficiently, increasing the overall amount of resources available to the pair at that time step. For example, if there are two possible resources, A and B, and the host and symbiont can choose only one resource type each, the host might choose resource A and the symbiont might choose resource B and enable the host to efficiently digest resource B. A host that overlaps with its symbiont is able to efficiently digest only a single type of resource and therefore fail to obtain half of the available resources.

The host first efficiently digests all of its available specialty resource. Its resource behavior value then determines the proportion of non-specialty resource types it gives to its symbiont. The symbiont then processes whatever types it can and returns a proportion of those resources to the host. The host is able to efficiently make use of the symbiont’s specialty resource after the symbiont processes it.

When the relationship is antagonistic and resource types are enabled, as shown in Figure 10, the chosen resource types determine the type of defense the host invests in and the type of attack the symbiont uses. By default 25 (example uses 10 for clarity) of each type of resource is again available to the host, but the symbiont is trying to steal some of them and the host is only able to defend one by default.

**Figure 10:**
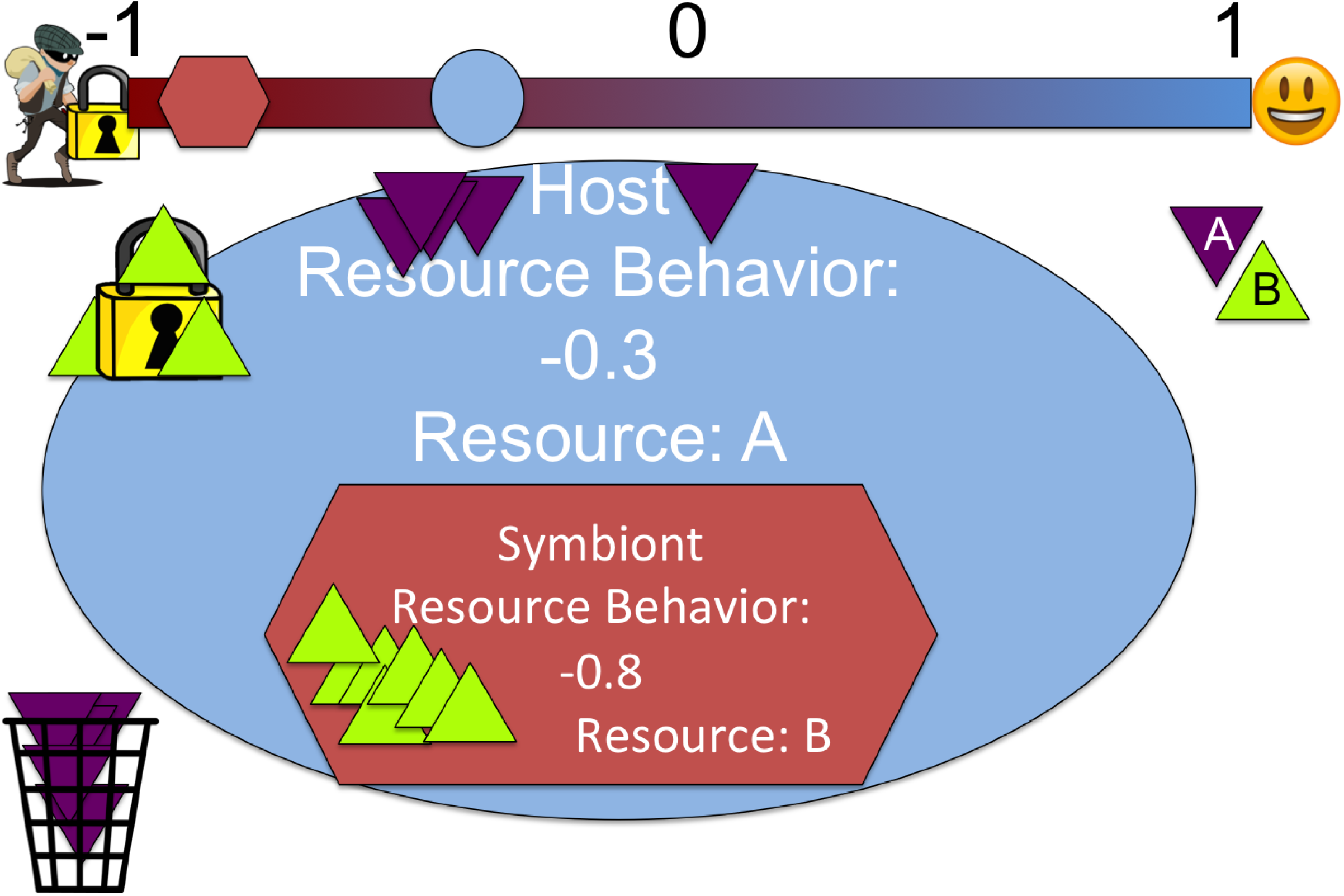
Example host and symbiont antagonistic resource behavior with non-matching chosen resource types. The host uses up 3 B in defense of resource A. The symbiont is able to destroy some of A and steal what is left of B.

If the host and symbiont do not have matching chosen resources, the host uses a portion – determined by its resource behavior value – of its non-specialty resources, in the example resource B, to protect its specialty resource. If the symbiont has a more negative resource behavior value than the host, it is able to overcome the host’s defenses and steal a portion of the host’s specialty resource in addition to what remains of the host’s non-specialty resources. If the symbiont has the same specialty resource as the host, the symbiont is only able to access that specialty resource, and the non-specialty resource is wasted. If the host is able to fend off the symbiont, each partner receives half of the B resource (host’s non-specialty), and the host keeps all of its specialty resource.

To summarize, when the host and symbiont are mutualistic, jointly they are more fit when they have different chosen resource types. However, when the relationship is antagonistic, the hosts will be most fit if they choose the *same* resource type as the symbiont. Conversely, the symbionts will be most fit if they choose a *different* resource type from the host, leading to conflict between hosts and parasites.

All combinations of resource behavior value and chosen resource type are presented in Table 2. These interaction dynamics were selected because they produce a fitness landscape that creates conflict between the partners. As seen in Figure 11, the region of the fitness landscape that is best for the symbiont is the opposite of that that is ideal for the host. This conflict is an abstraction of the many mutualism-parasitism systems where it is assumed that there is this conflict due to observed interactions.

**Table 2:** Interaction results for host and symbiont phenotypes with resource types

| Host Phenotype | Symbiont Phenotype | Chosen Resource Types | Result |
| --- | --- | --- | --- |
| $X > 0$ | $Y > 0$ | Types match<br>(ex: both A) | Host digests all of A, donates X of B to symbiont, inefficiently digests 1-X of B; symbiont cannot digest B and so X of B is wasted |
| $X > 0$ | $Y > 0$ | Types mismatch<br>(ex: Host A, Sym B) | Host digests A and donates X of B to symbiont, host inefficiently digests 1-X of B; symbiont processes B and returns Y proportion processed B, symbiont gains 1-Y of processed B; host digests processed B |
| $X < 0$ | $Y < 0$ | Types match<br>(ex: both A) | Host spends -X of B for general defense; symbiont tries to steal A, if $Y < X$, symbiont wastes Y-X of A and steals -Y of remaining A; if $Y \geq X$ each get half of A; B is wasted |

Table 2 – (cont'd)
| Host Phenotype | Symbiont Phenotype | Chosen Resource Types | Result |
| --- | --- | --- | --- |
| $X < 0$ | $Y < 0$ | Types mismatch (ex: Host A, Sym B) | Host spends $X$ of B for general defense; symbionts tries to steal B, if $Y < X$, symbiont wastes $Y-X$ of A, steals $-Y$ of B; if $Y \geq X$ each get half of B; host gets remaining A |
| $X > 0$ | $Y < 0$ | Types match (ex: both A) | Host donates $X$ of A to symbiont, symbiont takes all remaining A, B is wasted |
| $X > 0$ | $Y < 0$ | Types mismatch (ex: Host A, Sym B) | Host donates $X$ of A to symbiont, symbiont takes all of B, host gains what remains of A |
| $X < 0$ | $Y > 0$ | Types match (ex: both A) | Host digests A; Host spends $X$ of B on defense; Symbiont tries to digest A but none left; B is wasted |
| $X < 0$ | $Y > 0$ | Types mismatch (ex: Host A, Sym B) | Host digests A; Host spends $X$ of B on defense, another 0.5 of B is lost due to symbiont having to fight to get to it; Symbiont donates $Y$ of what B it gets to host as processed and keeps $1-Y$ of the B it got for itself |

**Figure 11:**
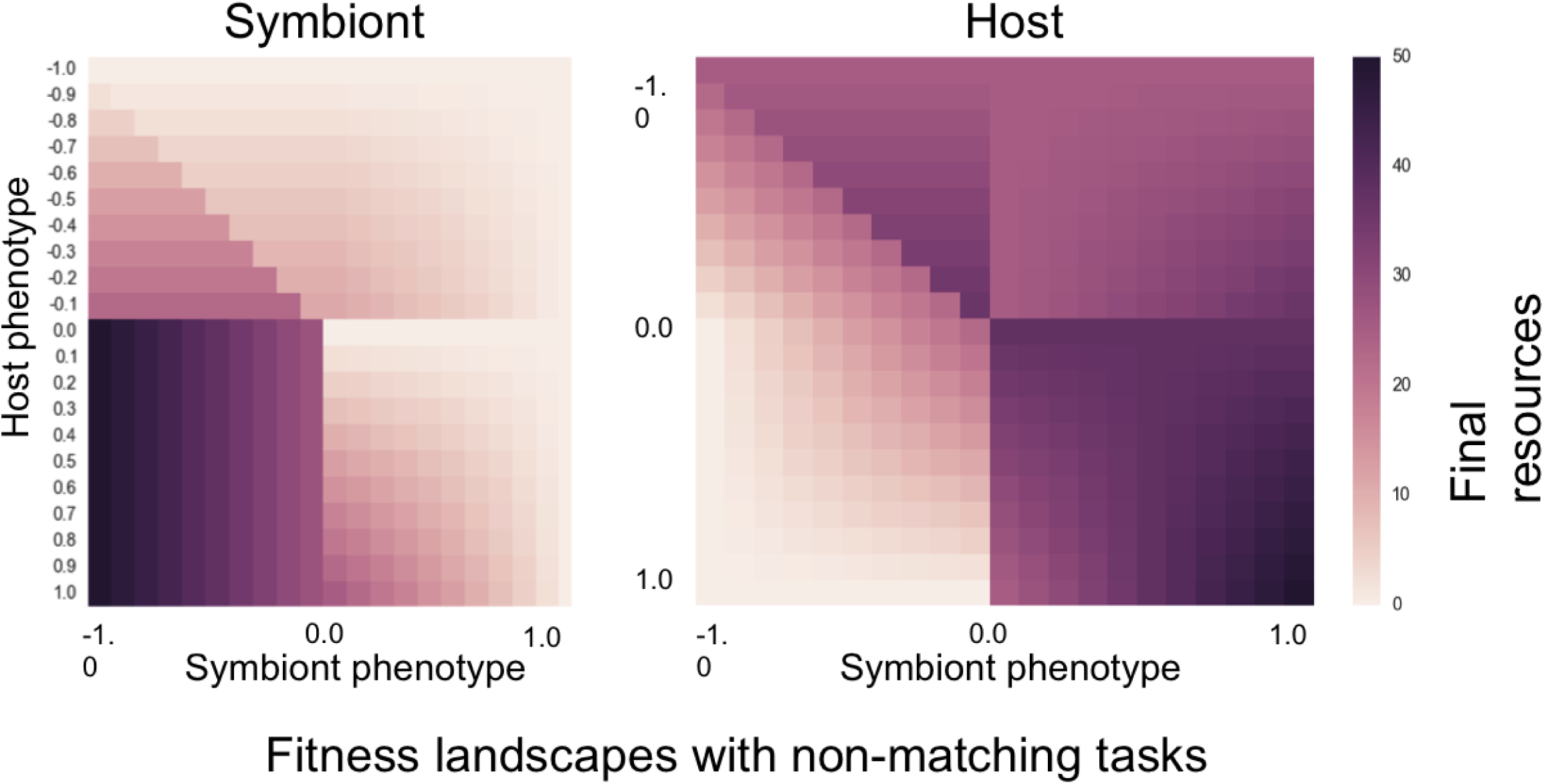
Fitness landscapes for Symbiont and Host with non-matching resource types. The symbiont and host fitness landscapes show that the two partners conflict on which phenotypes are best for each of them.

### 6.2 Is a mutualism able to evolve without a forced synergy effect?

By having the organisms choose resources to use, we are no longer enforcing an artificial synergy by multiplying the returned symbiont’s resources by 5. Instead, the host and symbiont only have a synergy if they use different resources. We first verified that the full range of parasitism to mutualism was able to evolve across vertical transmission rates, as seen in Fig. 12

**Figure 12:**
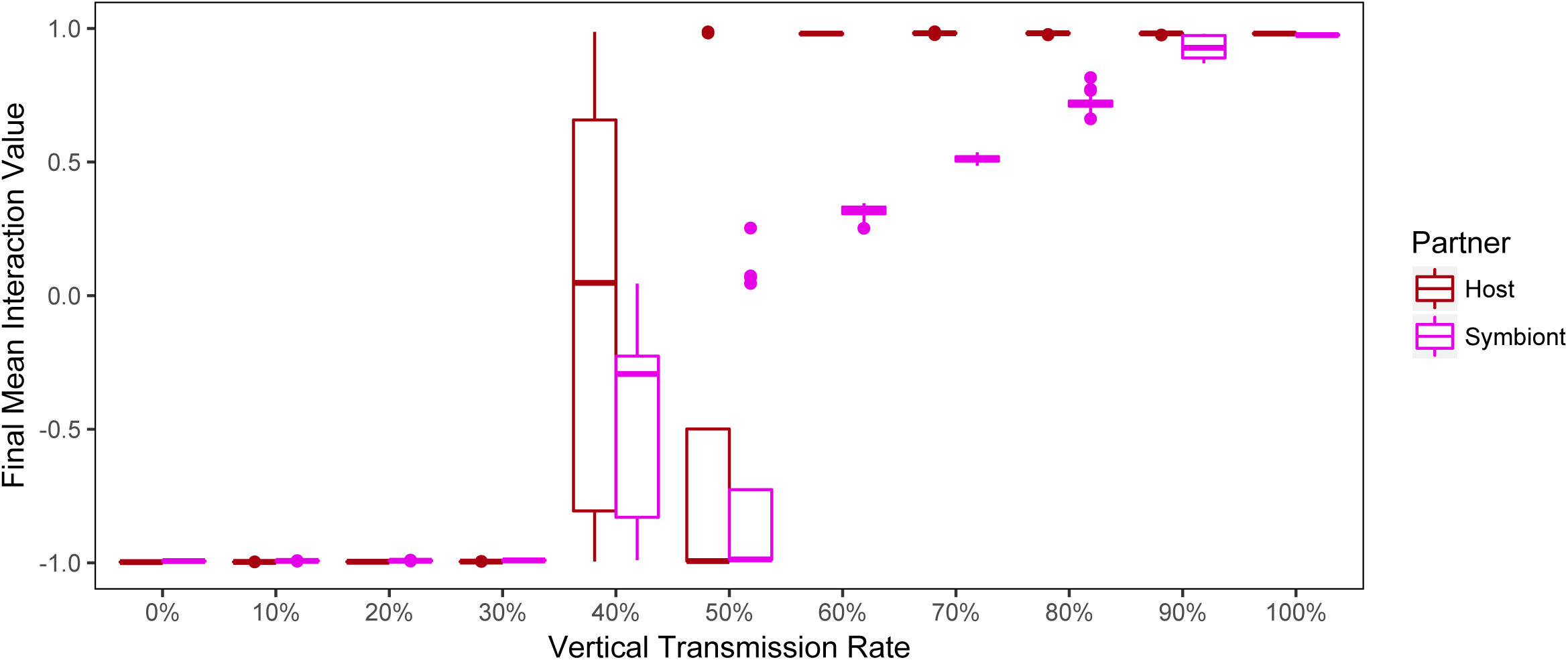
Evolution of mutualism across vertical transmission rates without artificial synergy. When an artificial synergy factor is replaced with a set of resource types that are divided based on the host and symbiont phenotypes, mutualisms are still able to evolve at higher vertical transmission rates.

We then determined that the coexistence of multiple symbiont phenotypes was not an artifact of the artificial synergy, as shown in Fig. 13.

**Figure 13:**
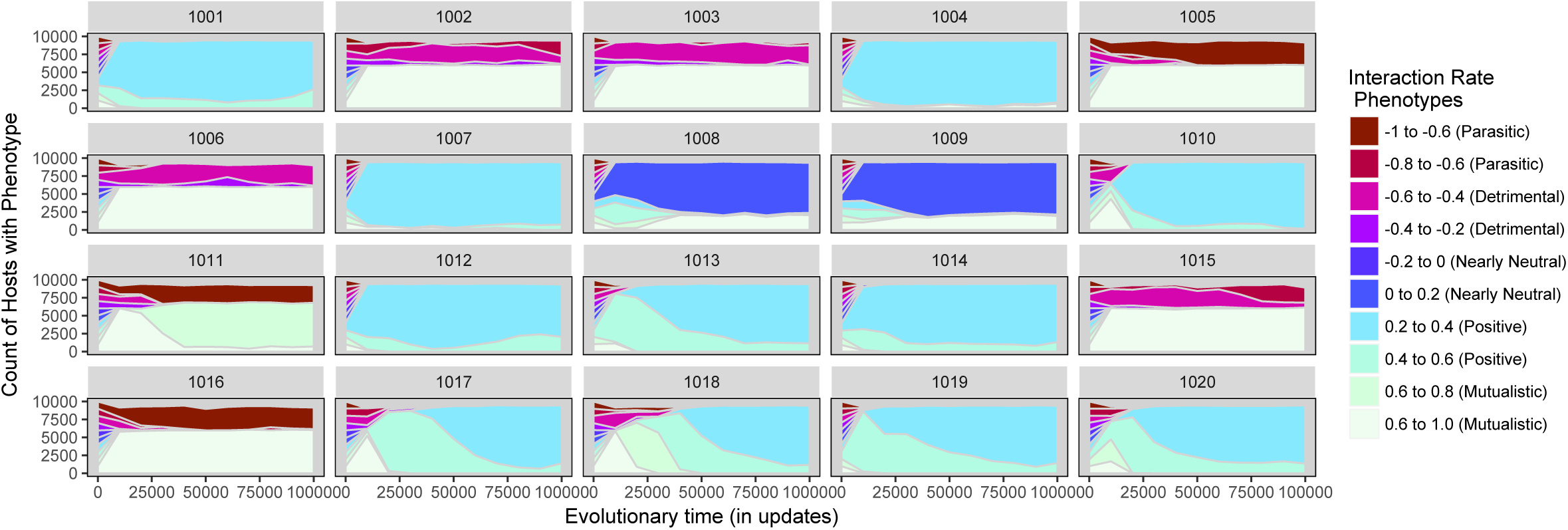
Count of symbiont phenotypes across evolution of mutualism when vertical transmission is 60%. As with a forced synergy, multiple phenotypes of symbiont, some parasitic and some mutualistic, are able to stably coexist with a more natural resource type system.

Further, the host population at vertical transmission rate 60% has multiple coexisting phenotypes, some defensive and some mutualistic, as shown in Fig. 14.

**Figure 14:**
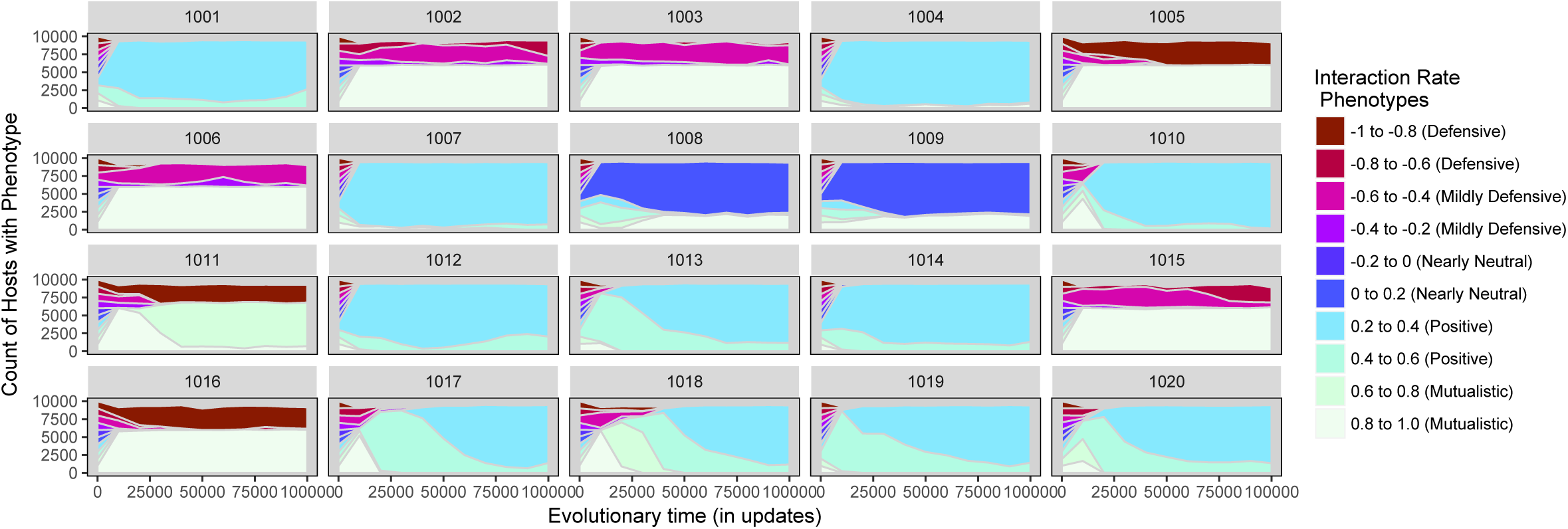
Count of host phenotypes across evolution of mutualism when vertical transmission rate is 60%. Host populations evolved coexistence of phenotypes.

Finally, as seen in Fig. 15, the resource type system enables the host to attempt to defend itself from parasitic symbionts by changing resource types. This change would then pressure the symbiont population to evolve to again match the host resource type, then pushing the hosts back to the original resource type. This dynamic is found in many natural systems of parasites and hosts [11], and therefore the emergence of this dynamic in Symbulation indicates that the Red Queen dynamic can occur in the simplest of host-parasite interactions.

**Figure 15:**
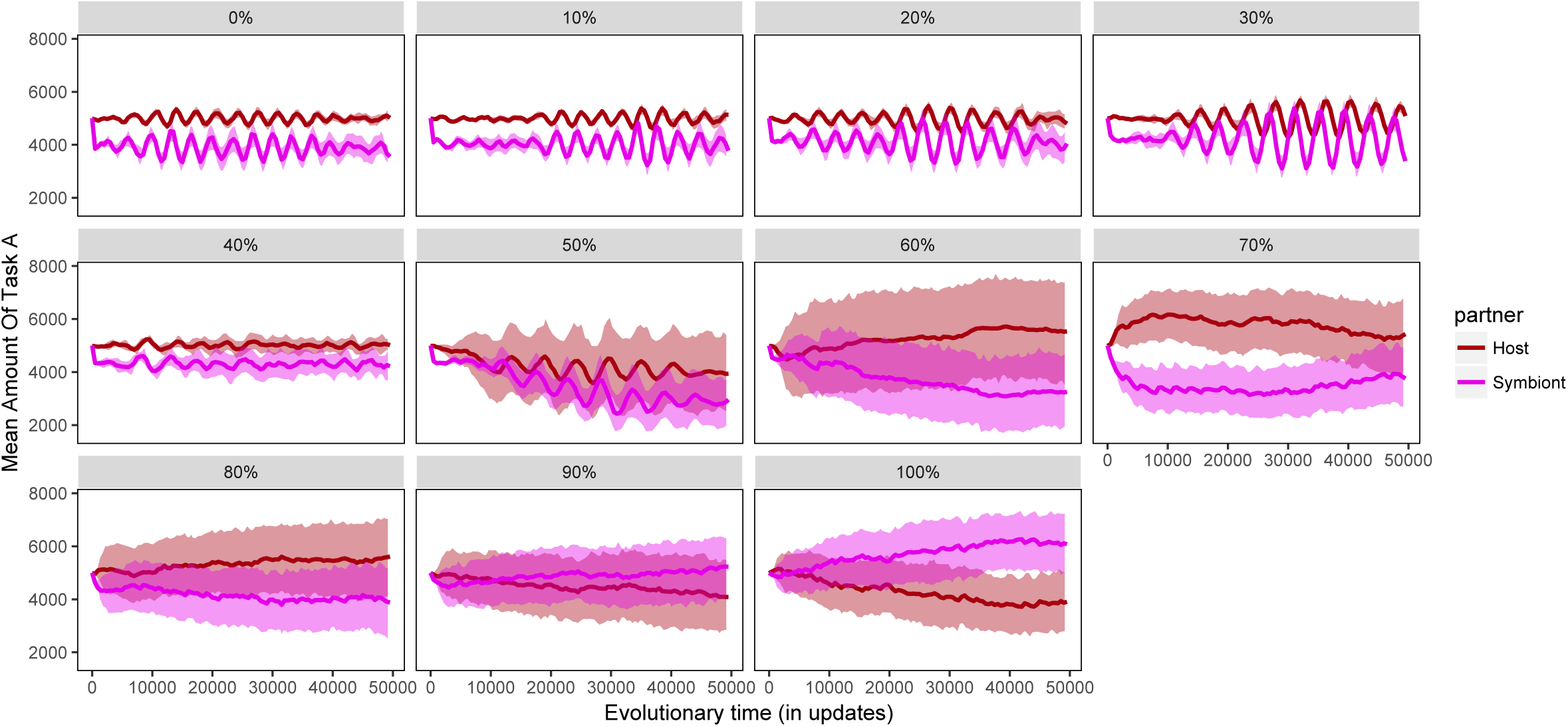
The number of hosts and symbionts choosing resource A over time across vertical transmission rates. When the vertical transmission rate is low and the majority of symbionts are parasitic, Red Queen oscillations emerge quickly in the populations.

### 6.3 Is division of labor between the partners necessary for the evolution of mutualism?

While we have shown that a mutualism can evolve and persist with a natural synergy created by a required division of labor, that does not mean that a division of labor is necessary for the evolution of a mutualism. To determine its necessity, we removed the limitation on how many resource types the host could choose, thus allowing the host to evolve to choose both resources A and B and no longer requiring a division of labor between host and symbiont.

When the host population is able to choose both resources A and B, it quickly evolves resistance to the symbiont, as seen in Fig. 16. The symbiont population in fact goes extinct, as seen in Fig. 17.

**Figure 16:**
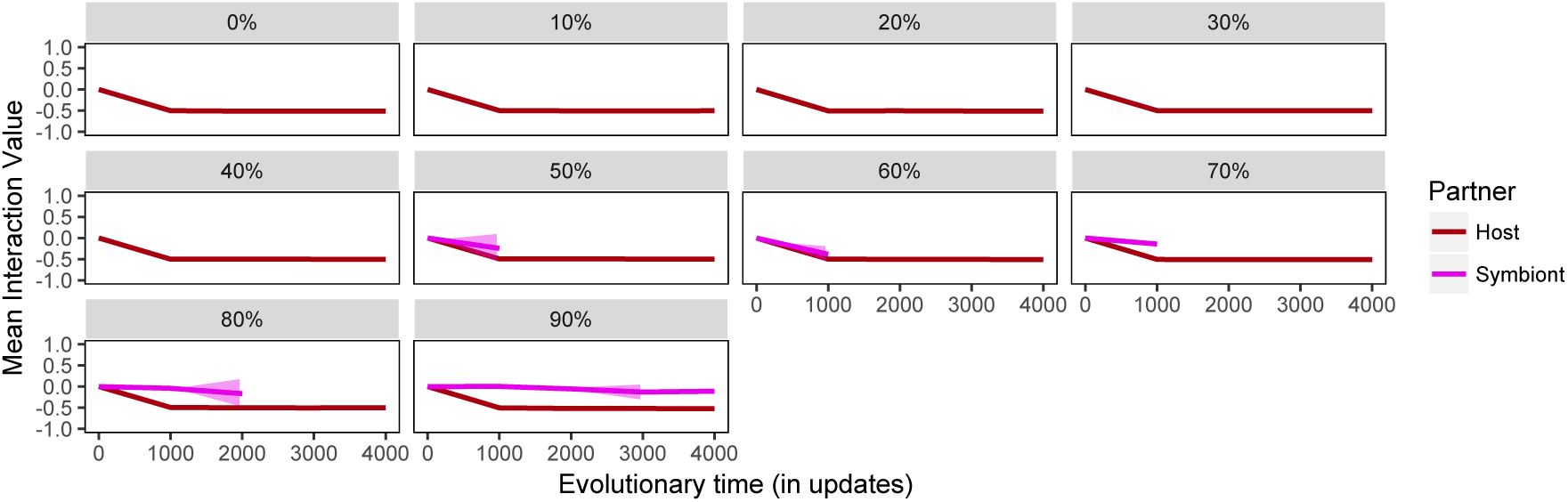
The resource behavior values of hosts and symbionts over time when the host can choose both resource types. The host quickly evolves to be extremely defensive and the symbiont population cannot evolve to parasitic before going extinct.

**Figure 17:**
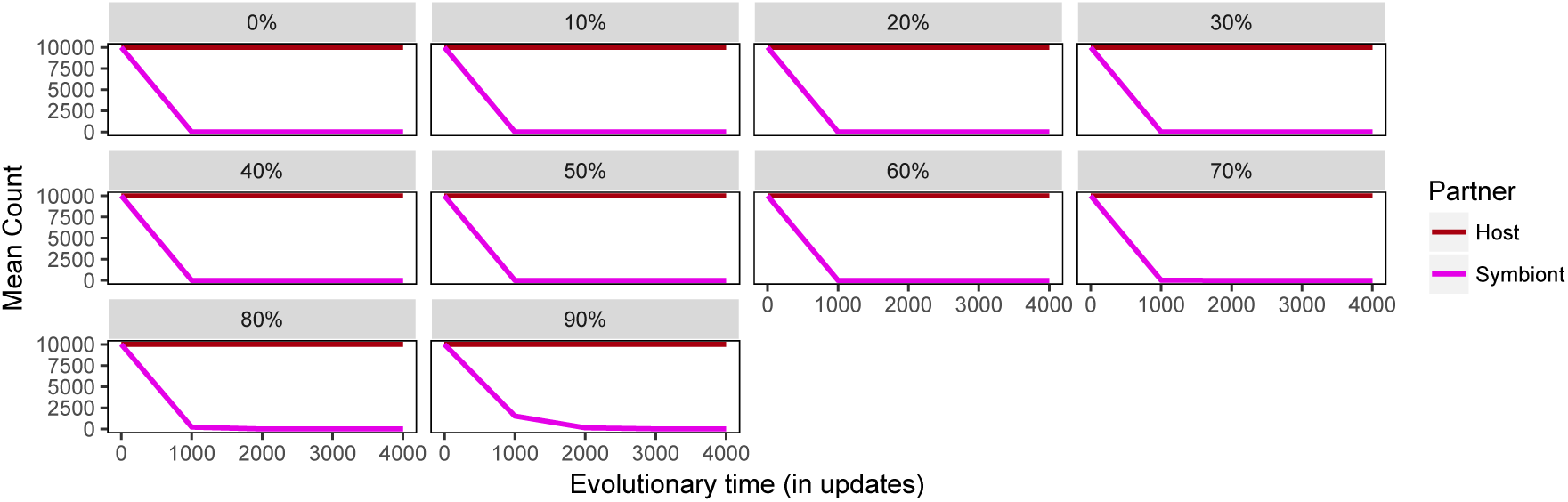
The population counts of hosts and symbionts over time when the host can choose both resource types. The symbiont quickly goes extinct at all vertical transmission rates except 100% (which was omitted because the host cannot force the symbiont extinct then).

This result indicates that some form of division of labor is necessary for the evolution of a mutualism, thus all of the following experiments have the required division-of-labor mechanism active.

## 7 Spatial Structure

Spatial structure is often able to increase the fitness advantage of cooperators because it allows them to preferentially interact with other cooperators and exclude cheaters [12]. Therefore, the introduction of spatial structure to a mutualism system has been hypothesized to increase the evolution of mutualism [33]. However, Akcay et al. presented a mathematical model where spatial structure in fact selected for less mutualism [2]. This result was due to the decreased diversity of symbionts in a given area, due to neighborhood dispersal of the symbionts. Akcay et al.’s system allowed hosts to be choosy regarding their symbiont partners and therefore a reduced diversity of symbionts to choose from meant that hosts in some areas were stuck with lower quality choices than the population as a whole. Conversely, in a well-mixed population, all hosts have access to the highest quality symbiont partners, potentially leading to more stable mutualism. Of course, if choosiness can evolve as well, as discussed in [16], consistently high quality partners can lead to reduced selection for choosiness. Mathematical models are limited in their ability to show how complex dynamics may combine in noisy and imperfect environments, making Symbulation an ideal system for exploring the effects of spatial structure on mutualism empirically.

### 7.1 Methods

For these experiments, we enabled neighborhood reproduction in Symbulation with two infinite resources (as discussed previously). When a host was able to reproduce, its offspring was randomly placed in one of the eight spaces surrounding the parent in a Moore neighborhood. Any previous occupant of the chosen space is killed, just as before in the well-mixed reproductive model. Symbionts are able to vertically transmit with these offspring with a set probability just as previously. If a symbiont is able to horizontally transmit, however, its offspring attempts to inject into one of the eight neighboring spaces. If there is not an uninfected host in the chosen space, the symbiont transmission fails, just as previously in the well-mixed model.

There is not an explicit mechanism for host choice in these experiments. Once a host has a symbiont, it cannot oust the symbiont. However, when a host is overwritten by another’s offspring, that offspring may be born uninfected. If that offspring is fairly similar to the organism in that space previously, the dynamic is nearly indistinguishable from a host ousting its current symbiont. A host with a less cooperative symbiont is more likely to be overwritten by a reproducing neighbor, thus making it more likely that a parasite will be ousted from its current host and replaced with an uninfected host. Conversely, a host with a more cooperative symbiont is more likely to produce offspring that have some probability of inheriting the host parent’s symbiont. Thus if a neighboring host is overwritten by a new host infected by a vertically transmitted symbiont, this dynamic approximates a host choosing a more mutualistic symbiont. I will refer to this dynamic as *implicit partner choice* as it closely parallels within-lifetime partner choice if host mutation rate is low enough.

### 7.2 How does local reproduction change the evolution of mutualism across vertical transmission rates?

Spatial structure or local reproduction is generally found to increase cooperation in single-species systems [12]. However, there is evidence that this might not be the case when a second species is included due to the limited number of partner choices available in a spatially structured environment [2]. Vertical transmission rate also affects the potential diversity of local organisms because the Symbulation system assumes that an established symbiont (whether passed vertically or horizontally) cannot be forced out of its host. Therefore, a higher vertical transmission rate would increase the number of hosts born already infected, potentially halting the spread of a horizontally transmitted strain and thus reducing local partner diversity. To explore how vertical transmission rate and spatial structure interact to influence the evolution of mutualism, we enabled spatial structure and compared the resulting data to the previous section’s.

As seen in Fig. 18, at vertical transmission rates of 60% and 70%, the final average resource behavior value is significantly higher with global reproduction than with local reproduction (60%: local mean −0.218, global mean 0.663, MannWhitney U test *p* << 0.005; 70%: local mean 0.385, global mean 0.759, MannWhitney U test *p* << 0.005). Conversely, when the vertical transmission rate is 30%, the average resource behavior value is significantly lower with global reproduction than with local reproduction (local mean: −0.693, global: −0.993, Mann-Whitney U test *p* << 0.005). This result indicates that the vertical transmission rate is a key determining factor in whether spatial structure selects for or against mutualism.

**Figure 18:**
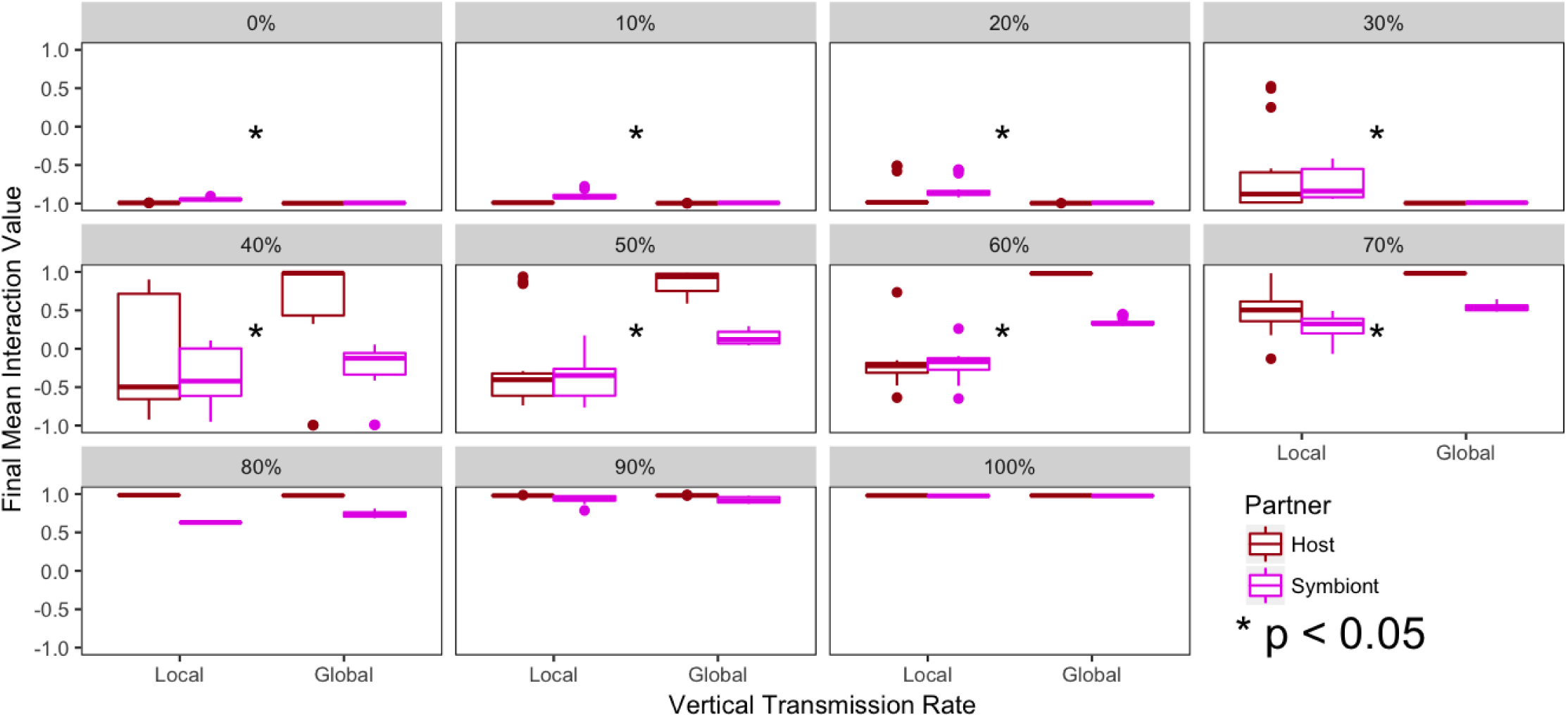
Resource behavior values after 100,000 updates across vertical transmission rates with global and local reproduction of host and symbiont. Considering the vertical transmission rates tested, at rates of 40% and lower, global reproduction resulted in significantly lower levels of mutualism than local reproduction. Conversely, at vertical transmission rates of 60% and 70%, global reproduction resulted in significantly less mutualism than local reproduction.

Akcay et al. predicted that spatial structure would reduce the fitness advantage for mutualisms because the hosts would not have access to the full population of symbionts when choosing partners [2]. While our experiments do not have an explicit partner choice mechanism – a host cannot oust a symbiont even if it is parasitic – this dynamic is approximated by population dynamics: a host with a low-quality partner is more likely to be overwritten before it has the chance to reproduce and possibly vertically transmit its partner. When that unfortunate host is replaced, it is likely to be by either an uninfected host or a host infected with a vertically transmitted mutualistic symbiont (since hosts with mutualists are more likely to be able to reproduce). Thus the same dynamic of low-quality symbionts being less likely to be maintained in a host through partner choice can be approximated by differential survival.

To determine whether our results are due to the same mechanism as outlined by Akcay et al., we calculated the average Shannon diversity of symbionts in a local neighborhood in both treatments. When reproduction is local, the Shannon diversity of symbionts will capture how many different phenotypes of symbiont a host lineage may ‘choose’. When reproduction is global, the Shannon diversity of the immediate neighboring symbionts will be a proxy for the Shannon diversity of the full population of symbionts.

As seen in Fig. 19, when the vertical transmission rate is 60%, global reproduction results in a significantly higher Shannon diversity of local symbionts than local reproduction. However at a vertical transmission rate of 30%, there is not a significant difference, as seen in Fig. 20.

**Figure 19:**
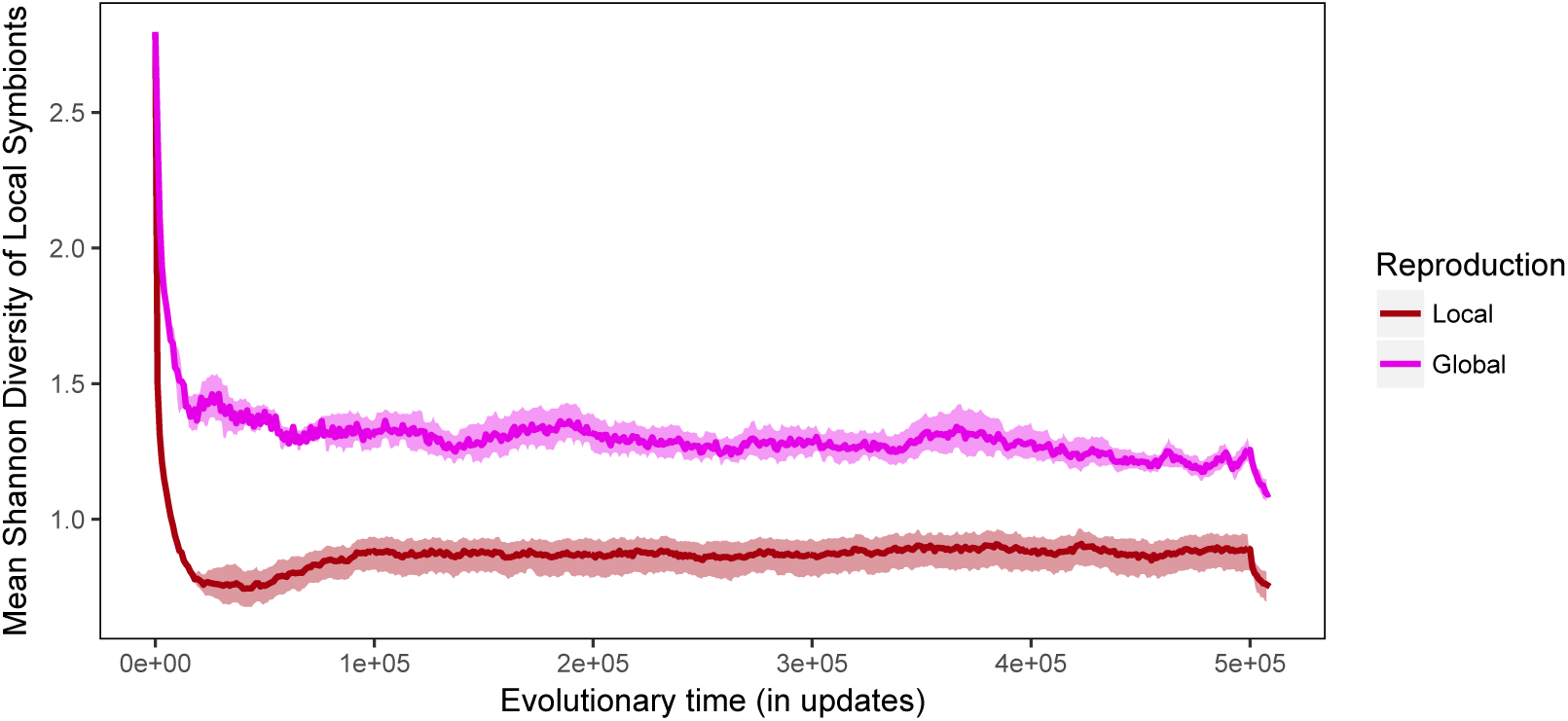
The average Shannon diversity of local symbionts when reproduction is local and global when vertical transmission rate is 60%. When reproduction is local, the Shannon diversity is lower than when reproduction is global, indicating the hosts have fewer choices of symbiont partner.

**Figure 20:**
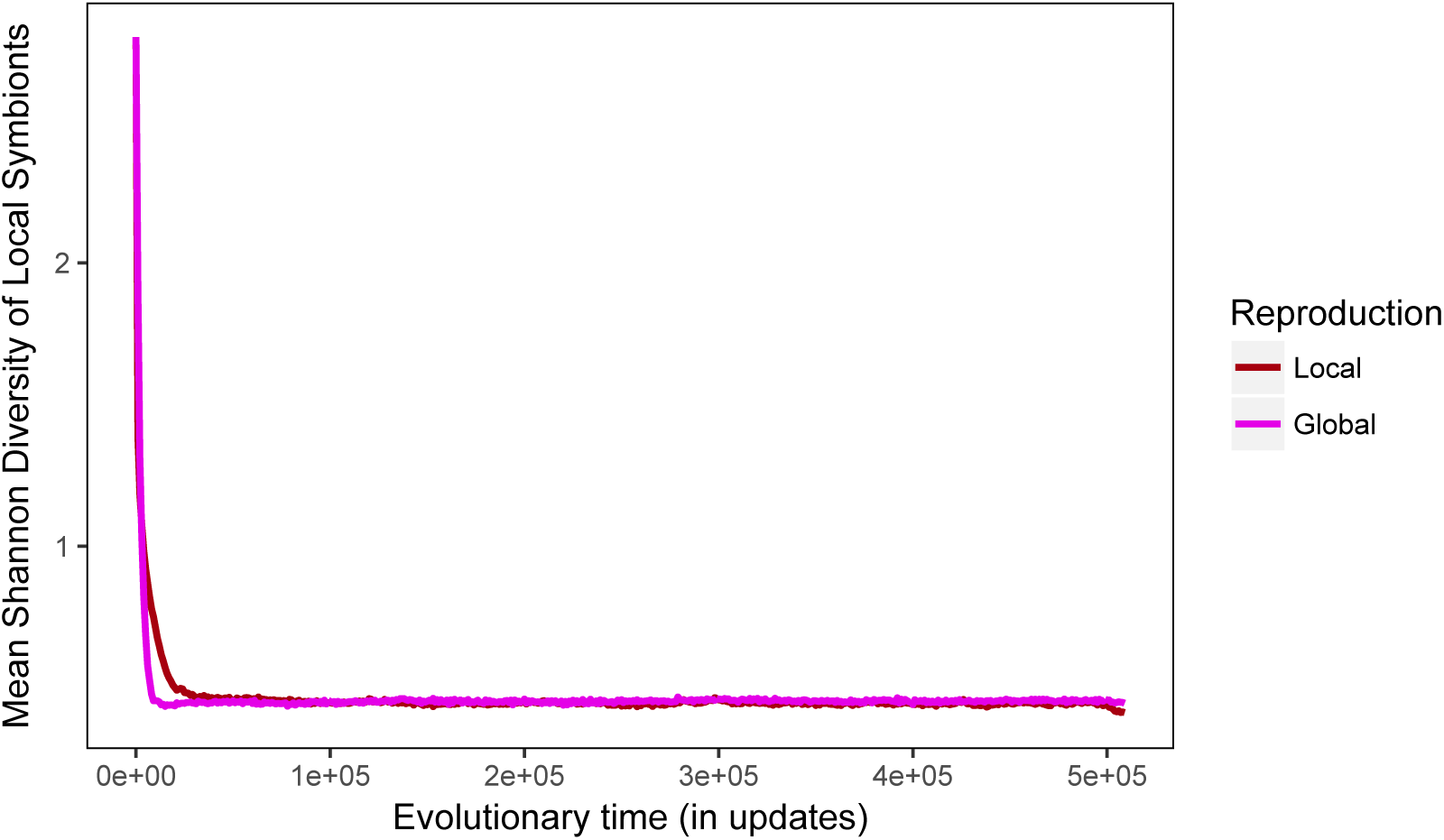
The average Shannon diversity of local symbionts when reproduction is local and global when vertical transmission rate is 30%. There is no significant difference between local and global reproduction.

This result confirms that spatial structure does reduce the partner choice options available to the host lineage at intermediate vertical transmission rates, leading to reduced evolution of mutualism. This result also shows that the the-oretical findings hold for a system with intermediate phenotypes and additional noise.

### 7.3 Does spatial structure increase the likelihood of mutualism evolving *de novo?*

Finally, we again return to the question of how the origin of a trait may differ from the maintenance of that trait. In the spatially structured environment, a mutualism that already exists in the population is able to invade in some vertical transmission rates, even if it is less than in a well-mixed environment. However, previously we showed that mutualisms were not able to evolve from neutral hosts and symbionts when we used an artificial synergy effect. The symbionts quickly died out and the hosts evolve defenses at all vertical transmission rates except 100%. While spatial structure reduces host partner choice, it also causes a newly emerged cooperative phenotype to stay next to its relatives, making it more likely a pocket of cooperation could grow to large enough numbers to become established. To determine the effect of spatial structure on the *de novo* evolution of mutualism, we again created a population of hosts and symbionts starting with neutral phenotypes, neither aggressive nor cooperative, and allowed evolution to proceed.

As shown in Fig. 21, the symbionts do not go extinct in the spatially-structured population, but a strong mutualism also does not evolve. At vertical transmission rates of 60% and below, parasites and defensive hosts become dominant by the end of the experiment. At higher vertical transmission rates, neutral or slightly positive hosts and symbionts are able to persist, though they do not evolve to be strong mutualists. This result suggests that, while a high vertical transmission rate is not sufficient for the *de novo* evolution of mutualism as it is for the maintenance, spatial structure does allow the symbionts to remain extant. This extra time could allow another mechanism to select for a mutualism, such as partner sanctions.

**Figure 21:**
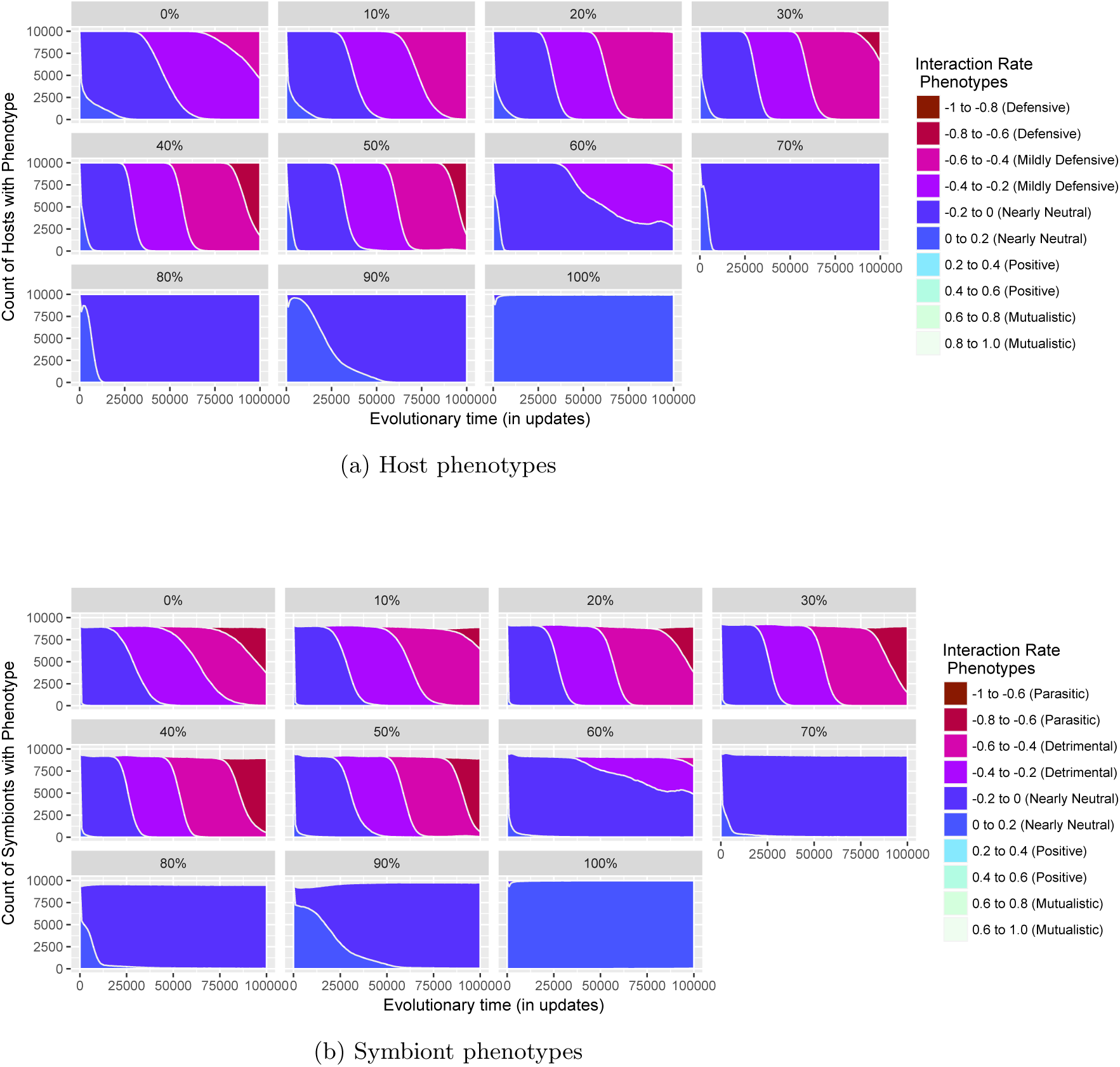
Host and symbiont phenotypes across vertical transmission rates when evolving from a neutral starting phenotype. At low vertical transmission rates, successive waves of more parasitic and defensive phenotypes evolve over time. At higher vertical transmission rates, neutral or slightly mutualistic phenotypes remain dominant over time.

## 8 Conclusion

This paper has presented the system Symbulation to study the many theoretical hypotheses related to the evolution of mutualism. Due to the limited nature of both biological systems and mathematical models, many of these hypotheses have been difficult if not impossible to test. Here, we have shown that a mutualism can evolve with imperfect vertical transmission and imperfect partners. We have empirically tested the theoretical hypothesis that the evolution of mutualism is in fact hampered by spatial structure at some vertical transmission rates due to lower partner choice. Finally, we have shown that the conditions under which a mutualism is able to increase in a population are not sufficient for the *de novo* evolution of mutualism in well-mixed or spatially-structured populations.

Taken together, these results argue the value of Symbulation as a new tool in the study of the evolution of mutualism that can be used to experimentally confirm theoretical findings and determine conditions that are likely to lead to the evolution of a mutualism in a natural system.

There are many expansions we plan to make to Symbulation to encompass more of the mutualism theory, including: within-lifetime partner choice, multi-infection, multi-births for symbionts, lysogeny, and partner sanctions. With these and other optional additional features, Symbulation will be a system that can be used to predict biological evolution of the human gut microbiome in response to changing conditions; pathogens of humans, crops, and livestock; and phage therapy in response to antibiotic resistant bacteria.

## 9 Acknowledgements

Thank you to Joshua Nahum for finding a key paper at just the right time. Thank you to the members of the DevoLab for feedback and proofreading.

This material is based in part upon work supported by the National Science Foundation under Cooperative Agreement No. DBI-0939454. Any opinions, findings, and conclusions or recommendations expressed in this material are those of the author(s) and do not necessarily reflect the views of the National Science Foundation.

